# Using splines for point spread function calibration at non-uniform depths in localization microscopy

**DOI:** 10.1101/2024.01.24.577007

**Authors:** Serafim Korovin, Dylan Kalisvaart, Shih-Te Hung, Jelmer Cnossen, Coen de Visser, Carlas S. Smith

## Abstract

Single-molecule localization microscopy methods extensively leverage the microscope point spread function (PSF) for fitting the molecules. Calibrating an accurate PSF model is especially difficult in the presence of depth-dependent aberrations which alter the PSF shape depending on the imaging depth. The aberrations at depths of a few micrometers become substantial enough to considerably impoverish the conventional calibration methods’ performance. In our work, we propose a novel spline model which enables the depth-dependent PSF model calibration by interpolating between the beads at arbitrary depths. We show that diffspline reduces the PSF intensity overestimation by 67.8 percentage points and underestimation by 21.8 percentage points. Moreover, it eliminates the depth-dependent bias and improves the localization precision two-fold compared to previous approaches.

## Introduction

Recent years have seen the emergence of single-molecule localization microscopy (SMLM)^1–3^ as a powerful tool for resolving three-dimensional (3D) biological structures^4–6^ beyond the diffraction limit. This technique relies on the sparse excitation of photoswitchable fluorophores to create isolated molecule images. The images contain molecule shapes blurred with the microscope emission pattern, or the point spread function (PSF), which can be experimentally designed and modelled to encode additional positional information in its shape^5,7,8^. The molecule positions are determined through maximum likelihood estimation (MLE)^9^ or maximum a posteriori (MAP) estimation^10^ of the position parameters within the point spread function (PSF) model. Therefore, creating a flexible PSF model that accurately represents the true experimental PSF is crucial for the ultimate localization accuracy and precision^11^.

To accurately model the PSF, piece-wise polynomials called spline functions^12^ were employed. Specifically, cubic splines (csplines)^13^, schematically depicted in Figure 1 (b), have been shown to enable the estimation precision that achieves the Cramèr-Rao lower bound (CRLB)^14,15^, the theoretical bound on the highest possible localization precision of unbiased estimators. By deriving the PSF model directly from the experimental data, splines provide an accurate PSF representation for varying PSF shapes^16^.

**Figure 1.**
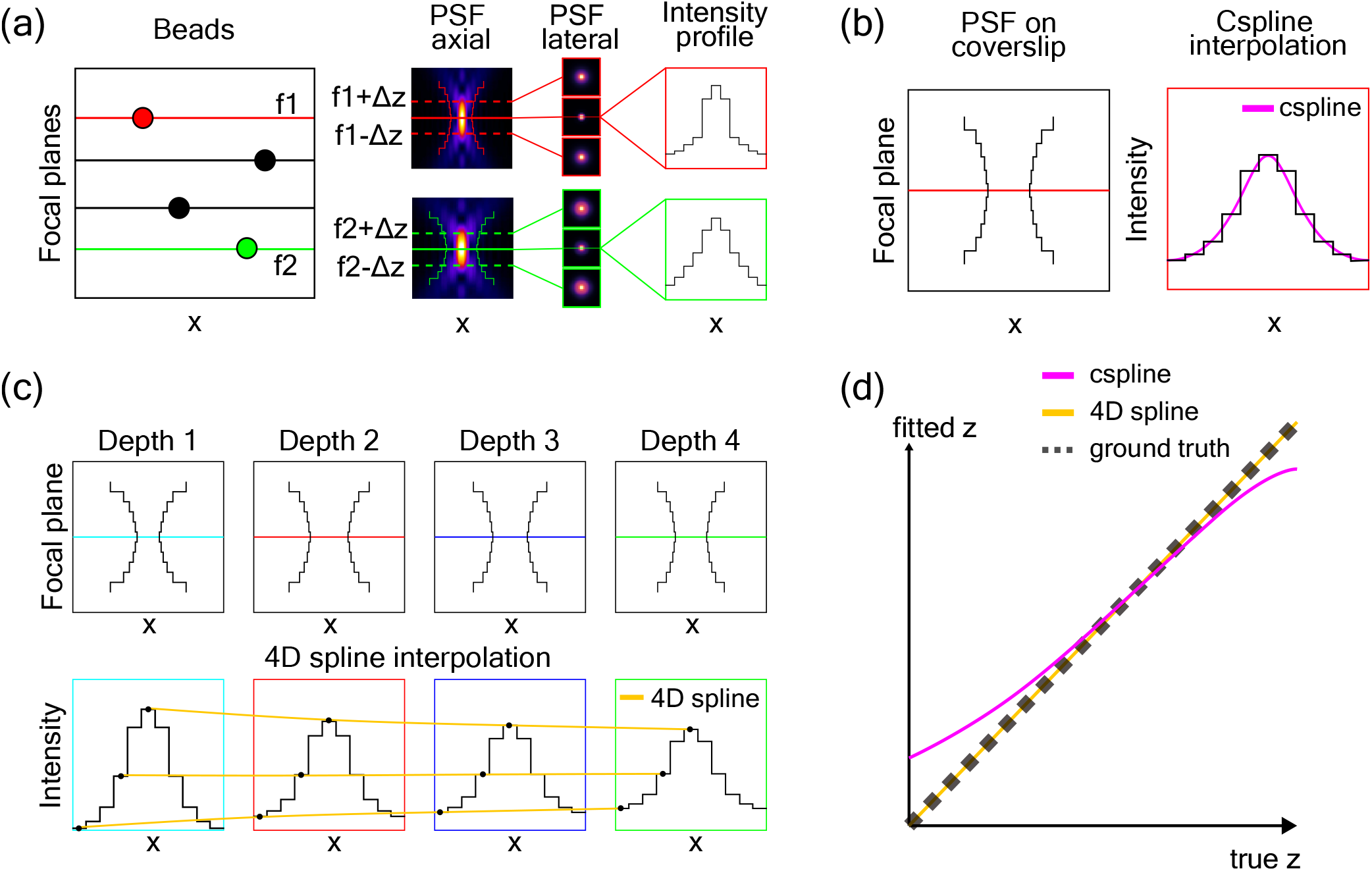
Schematic overview of this study. (a) Depending on the depth of the imaged beads, their 3D PSFs change. The leftmost figure depicts the beads located at various focal planes. Next to it, we show the axial and lateral views on the corresponding PSFs, with the PSF centers being located at the corresponding nominal focal planes. The colored outline of the PSF schematically represents the PSF shape obtained by an intensity cutoff. The rightmost figure shows the PSF intensity profiles in *ZX*, where the difference between the intensities at the given depths can be clearly observed. (b) 3D cspline approach creates a smooth PSF model by interpolating between the given intensity values. The 3D cspline model is calibrated on the bead data from the coverslip. (c) 4D spline can also interpolate between the bead data from different arbitrary depths. This way, we can obtain 3D PSF models from the depths that are not present in the data. (d) Depth-dependent 4D spline interpolation produces more accurate PSF models, and thus results in more accurate z-localizations.

Spline models are conventionally calibrated on image frames obtained by scanning through the bead sample along the *Z*-axis.The scanning is performed by moving the focal plane through the sample with a fixed step in *Z*. The bead PSFs can be obtained from those frames by taking the nominal focal plane as the center and collecting frames within a range of *Z*-positions around it^17^, which is depicted in Figure 1 (a). For the spline model calibration, the resulting *Z*-stack of images contains fluorescent beads which are usually located on the coverslip. However, when imaging deeper into the sample as illustrated in Figure 1 (a), depth-induced spherical aberrations emerge and become exacerbated in the presence of the refractive index mismatch between the objective lens immersion oil and sample medium^18^. These aberrations lead to considerable errors in PSF shapes, even for depths of only a few micrometers^18^.

Several methods have been proposed for addressing the challenges posed by depth-induced aberrations. Active aberration correction methods^19,20^ aim to estimate the aberrations directly from the wavefront, but these methods require complex adaptive optics hardware and are difficult to operate for non-experts. Other approaches use phase-retrieved PSF methods^21–25^ to construct a pupil function that describes the magnitude and the phase of the emission light wavefront. However, such methods require specific imaging system parameters (e.g., refractive indices and emission wavelengths) to build the pupil function, and thus are susceptible to parameter deviations in the form of noise or optical imperfections.

In turn, a cspline-based approach^17^ was devised to correct depth-induced aberrations without any requirements for optical parameters or hardware. It estimates the cspline bias for every depth and subtracts the bias from the localization estimates.

However, since the correction is only applied to the final estimates directly, the PSF model remains incorrect. This means that the localization precision is unaffected by the correction. Therefore, depth information must be incorporated into the spline PSF model for localization to be accurate and precise.

An alternative strategy is to extend the cspline model from three-dimensional (3D) to four-dimensional (4D) by interpolating between the point spread functions (PSFs) at various depths. However, this approach is computationally expensive, as the number of coefficients for which the spline has to be calibrated increases exponentially with the spline dimensionality. This issue was alleviated with the multivariate B-spline^26^ which employs basis functions to obtain the interpolation, thus reducing the coefficient complexity to depend only on the number of voxels present in the PSF image. However, applying B-splines to depth-dependent PSF calibration is challenging, since B-splines assume uniformly spaced calibration points in all dimensions, while the actual bead depths in the calibration *Z*-stack are not uniformly spread.

To this end, we propose a depth-dependent spline PSF model formulated as a non-uniform Catmull-Rom spline. Similar to the B-spline, the Catmull-Rom spline^27^ reduces the coefficient complexity as it builds a spline function based on pixel values. Furthermore, Barry and Goldman^28^ proposed a recursive algorithm that allowed the Catmull-Rom spline to be built on arbitrarily spaced points. By applying Barry and Goldman’s definition of the Catmull-Rom spline, we were able to build a spline PSF model capable of interpolating between beads at non-equidistant depths as illustrated in Figure 1 (c).

The resulting non-uniform Catmull-Rom spline, which we name Depth-Interpolating Fitting Function spline (diffspline), is evaluated against the cspline approach on simulated beads at various depths, as schematically shown in Figure 1 (d). We compared the diffspline with the cspline in terms of the PSF shapes and localization errors (bias, precision, and CRLB). We demonstrated that the diffspline PSF produces a more accurate PSF shape, with the intensity overestimation reduced by 67.8 percent points and with the underestimation reduced by 21.8 percent points. The effectiveness of the diffspline for deep sample localization was demonstrated by achieving an approximately two-fold improvement in localization precision compared to the coverslip-calibrated cspline. Moreover, the precision of our model closely followed its CRLB, suggesting that the diffspline delivers the correct CRLB for the evaluated depth.

## Data pipeline

We developed a simulated pipeline that generates PSF *Z*-stacks for any given depth, and we leveraged the pipeline to verify our approach. First, we describe the general data acquisition process for the experimental data. Then, we describe the implementation of the simulated dataset.

### Data acquisition and processing pipeline

The data acquisition is done in line with the previous work^13,17^. The fluorescent beads are immobilized on the coverslip at different depths. First, a 3D *Z*-stack of the bead sample is obtained by moving the focal plane with a fixed step size. The next step involves detecting the beads from the *Z*-stack. This can be done by maximum intensity projection and thresholding^13^. As a result, the regions in *X*,*Y*,*Z* which contain the beads are found. The depth values are typically separated into the nominal focal plane, which corresponds to the depth at the center of the bead, and the *Z*-positions, corresponding to the depths relative to the nominal focal plane of the bead. To find the nominal focal plan of each bead, an approach by Li et al.^17^ can be employed wherein a coverslip-calibrated cspline is fitted on the *Z*-stack, and the frame that returns *Z* = 0 is assigned as the nominal focal plane. To improve the resulting spline calibration fidelity from the noisy *Z*-stacks, several realizations of each bead are collected by shifting the stage in *X* and *Y* -directions and taking z-stacks at the shifted FOV, as depicted in Figure 2 (a, b). The resulting PSF realizations of the same bead are aligned and averaged, as represented in Figure 2 (c), which gives a better signal-to-noise ratio.

**Figure 2.**
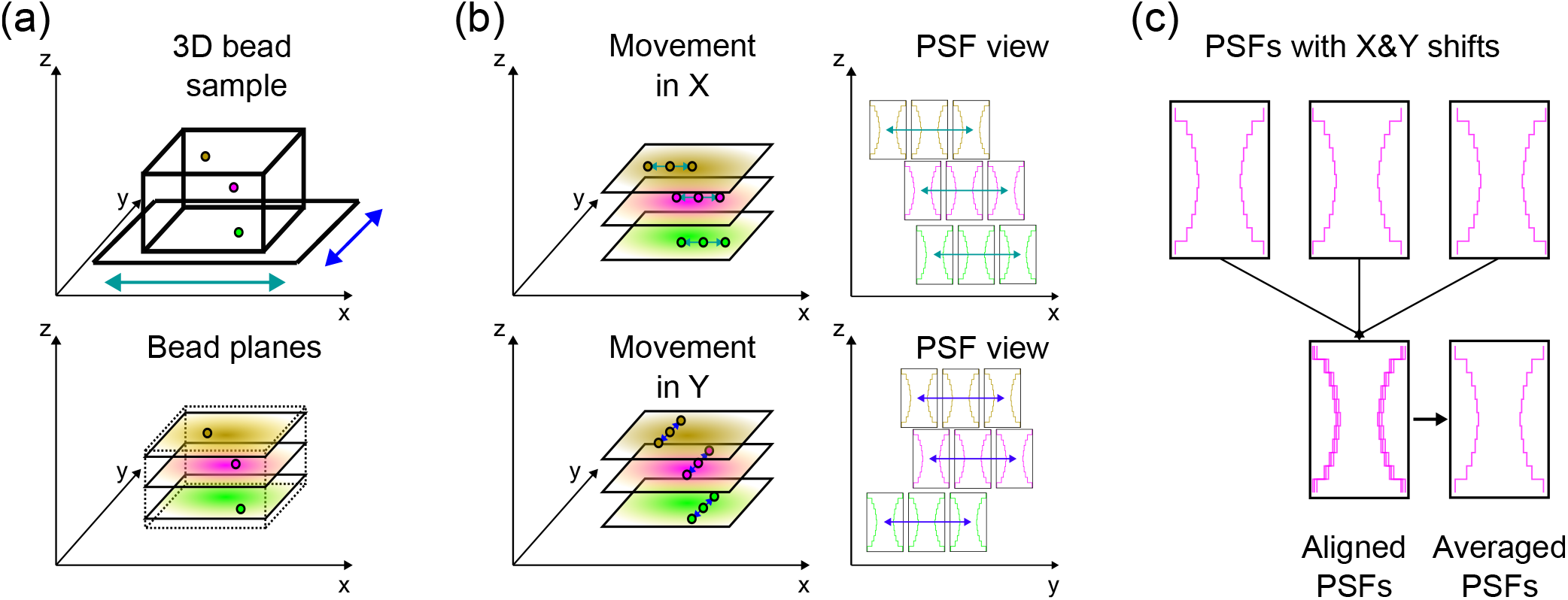
Overview of the processing pipeline that we simulate in our work. (a) The simulated sample contains beads at multiple depths. The sample stage movement along the *X* and *Y* axis is included in the simulation. (b) The simulated bead sample is moved in both *X* and *Y* planes and scanned at various positions. In this manner, we can acquire different realizations of the same bead across the field of view. (c) The obtained realizations are aligned and averaged into one PSF.

### Simulated pipeline

In the simulated pipeline, in order to measure the effects of depth in isolation from the detection error effects, we simulate the PSFs directly skipping the detection part of the original pipeline. A 3D bead sample is simulated by generating PSF *Z*-stacks at the given depths using a vectorial model^29^. The bead PSFs at different depths are simulated by adding first- and second-order spherical aberrations, as proposed by Booth et al.^18^. Each *Z*-stack contains one bead, which is assigned a random intensity between 5000 and 15000 photons, and a constant background of 50 photons/pixel. Moreover, we simulate the shot noise in the *Z*-stack with a Poisson distribution^9^.

The FOV shifting is simulated by adding a random position phase in the pupil phase function of the vectorial model. The shift is sampled from a uniform distribution ranging from 0 to the pixel size in *XY* .

In our simulations, we use the emission wavelength *λ* = 680 nm, an astigmatic PSF with vertical astigmatism 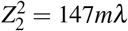, the refractive index for the sample medium is 1.33 and for the immersion oil is 1.52. The simulated voxel size is 100 nm in *X* and *Y* axes, and the *Z*-range spans from −500 nm to 500 nm with 10 nm step.

Subsequently, we apply the alignment and processing methods proposed by Li et al.^13^. First, all PSFs are gathered and aligned around a reference bead^13^. Next, the PSFs are divided into groups containing realizations of the same bead. Then, for each group, we apply the processing by Li et al.^13^ wherein we average the bead PSFs, eliminate the background by *Z*-stack minimum subtraction, normalize the PSF amplitude by the cumulative intensity of the central axial slice, and smooth the PSF shape in *Z*-direction with a smoothing B-spline. As an outcome of the processing, which is depicted in Figure 2 (c), we obtain a processed PSF *Z*-stack for each simulated bead. This *Z*-stack is the output of the simulation pipeline.

## Spline calibration

We calibrate spline models on the processed PSF *Z*-stacks. The cspline is calibrated on the simulated PSF *Z*-stack on the coverslip (0 nm depth)^13^.

For the diffspline calibration, we first generate simulated beads at depths *d*_*i*_, which are specified separately for each experiment. To calibrate the diffspline at a specific depth *d*_*eval*_, we interpolate between four consecutive PSFs with depths *d*_*i*_ closest to *d*_*eval*_, i.e. we find *i* s.t. *d*_*i*_ : *d*_*i*−1_ *< d*_*i*_ ≤ *d*_*eval*_ *< d*_*i*+1_ *< d*_*i*+2_). The interpolation is performed with the non-uniform Catmull-Rom spline, as described in Supplementary Note S1. We further discuss the effects of the Poisson noise on the spline calibration in Supplementary Note S1. Thus, we obtain an interpolated 3D PSF model for the required depth. Since csplines have an established localization methodology^16^, we convert the interpolated 3D Catmull-Rom spline into a cspline form for a direct comparison between the two spline types. The conversion is performed by computing the spline coefficients with Catmull-Rom and saving them in a cspline model.

Then, we evaluate the converted model by using it to localize on the simulated data. To do so, we first use the vectorial model to generate PSFs at the image center in *XY* for a given depth with an intensity of 10000 photons and a constant background of 50 photons/pixel. Next, we create 100 realizations of each *Z*-position by applying the shot noise to the PSF. Following that, we obtain the maximum likelihood estimates of the emitter *Z*-positions from the generated frames with the Levenberg-Marquardt algorithm^30^ using the damping factor = 10 and 100 iterations. Finally, we calculate the localization bias and precision from the estimates given the known ground truth. The bias is calculated as the mean error, and the precision is calculated as the standard deviation of the estimates over 100 frames. For the PSF shape evaluation, we employ the pixel-wise relative difference between the target PSF and the ground truth PSF, which is the normalized version of the pixel-wise difference used by Babcock et al.^16^. It is calculated as the difference between the target and ground truth, then divided by the ground truth pixel values. This metric, multiplied by 100, describes the percentage of deviation from the ground-truth reference.

## Results

Several experiments were conducted to quantify the diffspline performance. First, we compared it with the cspline approach by evaluating the splines on the beads that were present in the z-stack (“on-bead evaluation”). This tested the capabilities of the splines as PSF models without interpolation effects. We evaluated the spline PSF shapes and localization metrics, such as bias and precision. Furthermore, we tested the interpolation capabilities of the diffspline model. We calculated the diffspline localization bias and precision for depths between uniformly and nonuniformly spaced beads. Moreover, we investigated the influence of the distance between beads on the resulting localization errors.

### On-bead evaluation

First, we compared the cspline approach with our method by evaluating them on PSFs at 6000 nm depth. The simulation pipeline was leveraged to generate PSFs at 0 nm depth (coverslip), which was used for the cspline model calibration, and 5900, 6000, 6100, 6200 nm, which were used for the diffspline calibration. Both models were evaluated at depth *d* = 6000 nm.

Figure 3 shows that the 4D diffspline was able to capture the PSF shape more accurately than the cspline, which had significant shape deformations when compared to the ground truth (GT). The largest negative relative difference, or under- estimation, in the pixel-wise relative intensity difference between the cspline and the GT being −63.5% at *Z* = −500, and the largest positive difference, or overestimation, is 76.3% at *Z* = 500 (see Supplementary Note S1). In contrast, the largest under- and overestimation for the diffspline were −41.7% at *Z* = 500 and 8.5% at *Z* = −500. Therefore, we observed that the largest intensity overestimation was reduced by 67.8 percent points, whereas the largest underestimations were reduced by 21.8 percent points. Moreover, as can be observed in Figure 3, the diffspline underestimations followed the general PSF shape, and thus could be attributed to an insufficient shape amplitude rather than an incorrect lateral shape profile.

**Figure 3.**
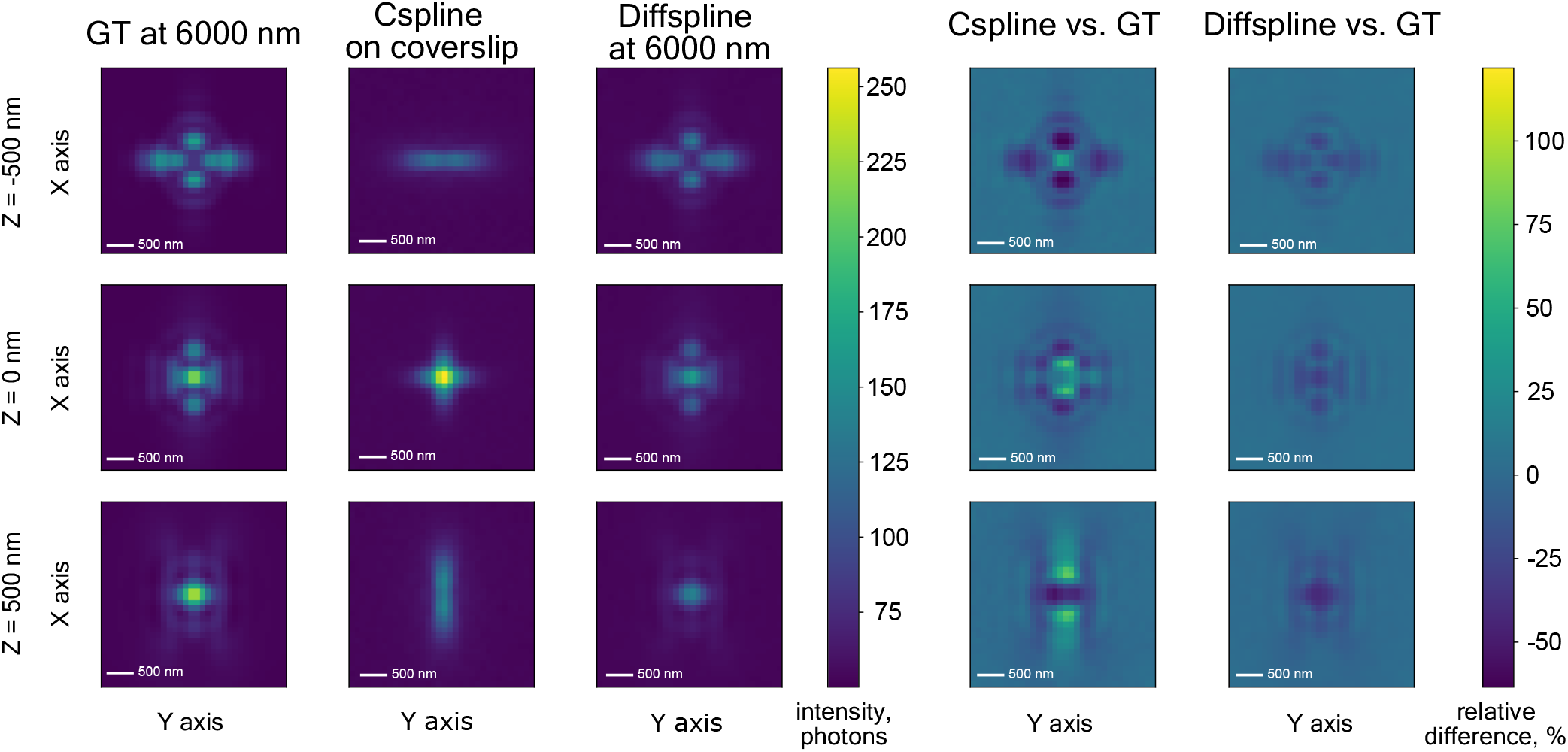
PSF visualization - comparing cspline and diffspline PSFs to the simulated ground truth (GT) at the 6000 nm depth for the bead intensity = 10000 photons and the background of 50 photons/pixel. The first three columns show the PSF intensity in *X* and *Y*, and the last two columns show the respective pixel-wise difference from the ground truth in percentages. Since the cspline is calibrated on the coverslip, it fails to capture the depth-dependent aberrations at larger depths and therefore considerable shape differences with the simulated ground truth occur.

For the estimation results, the coverslip-calibrated cspline at the depth of 6000 nm displayed a large bias, as can be observed in Figure 4(a). Although aberration-correction^17^ can be employed to correct the depth-induced bias in the cspline approach, the precision remains unaddressed, which results in highly imprecise localizations, as shown in Figure 4(b). In contrast, the diffspline approach displays a consistently low bias of approximately −5 nm on average. Moreover, Figure 4(b) demonstrates that the diffspline precision remains consistently low at approximately 12 nm on average, while it stays close to the CRLB with the difference not exceeding 4 nm.

**Figure 4.**
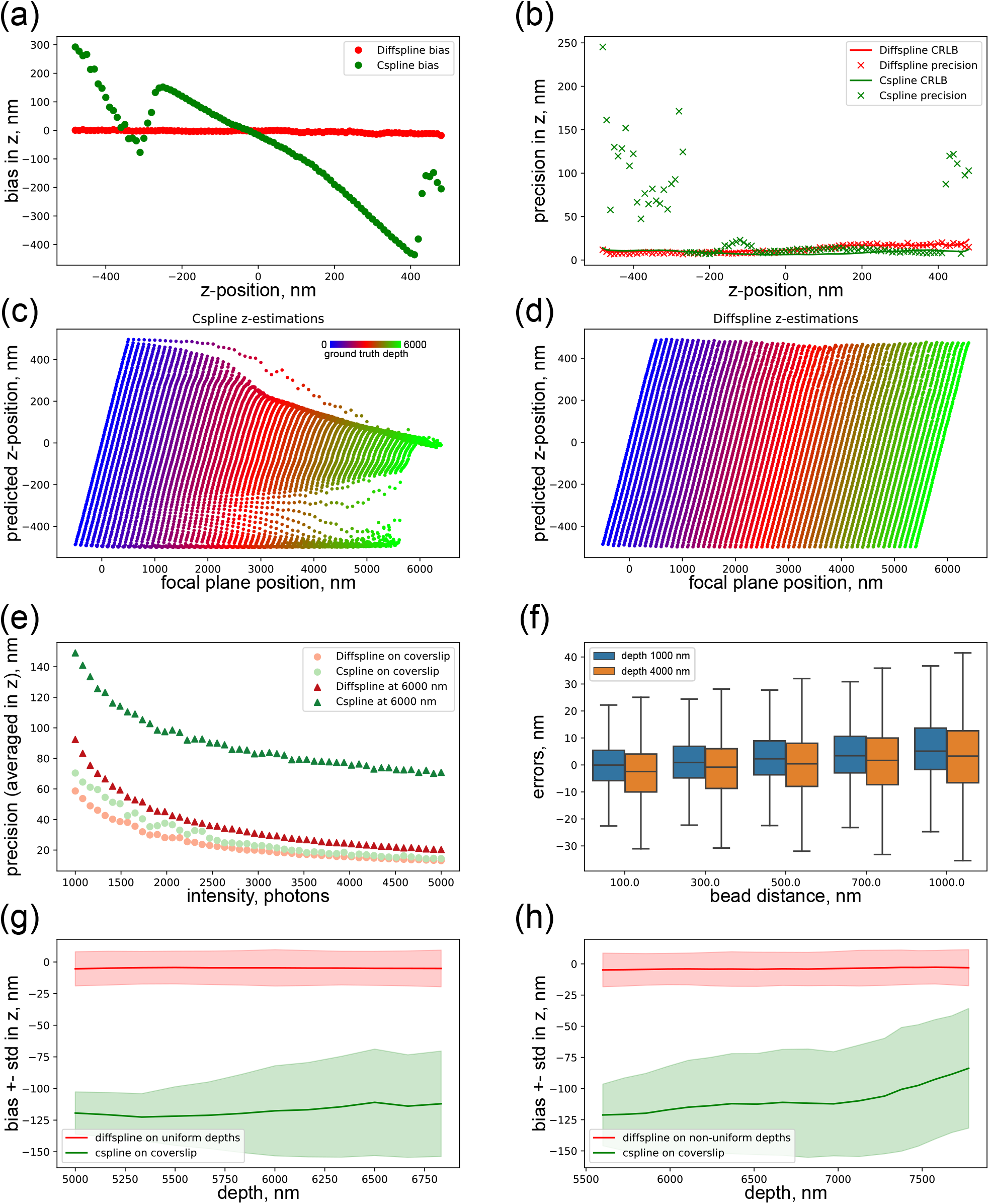
Diffspline localization performance quantification. (a, b) We compare the coverslip-calibrated cspline and diffspline approaches at 6000 nm depth by measuring their localization biases (average errors) and precisions (standard deviations of errors) with CRLBs (theoretical lower bound of the standard deviations). (c) Estimations produced by the cspline model calibrated on the coverslip. (d) Estimations by the diffspline calibrated on the simulated deep bead sample. (e) Diffspline and cspline precisions were compared over varying intensities at different depths. (f) Diffspline localization error distributions are plotted for different distances between the calibration beads at 1000 nm and 4000 nm depths. (g, h) Evaluating the diffspline interpolation between the uniformly (g) and non-uniformly (h) distributed beads against the cspline approach.

Furthermore, we demonstrate the precision advantages of our method by comparing cspline and diffspline precisions. We evaluated the cspline and diffspline models on frames with bead intensities varying from 1000 to 5000 photons with a constant background of 50 photons/pixel. The evaluation was performed by calculating the precision averaged over the *Z*-range. Figure 4(e) shows that the diffspline displays lower precision for all depths and intensities than the cspline. The diffspline precision is lower by 60 nm on average when evaluated at a 6000 nm depth with a 1000 photon intensity and by approximately 45 nm on average with 5000 photon intensity. This precision difference is lower when evaluated on the coverslip, where the difference of 10 nm at 1000 photon intensity is almost eliminated with an increase in intensity to 5000 photons. The lack of a similar decrease for the 6000 nm depth suggests that the model mismatch between the real PSF and the cspline prevents the latter from localizing more precisely.

To provide a more detailed look into the distribution of localization estimates over depth, we evaluated cspline and diffspline models at every depth from 0 to 6000 nm with a 100 nm step. For every depth, we estimated *Z*-positions for *Z*-range from −500 to 500 nm with a 10 nm step for 100 simulated frames, and averaged the predictions over the frames. The resulting *Z*-position estimates were plotted against the ground truth depths, as depicted in Figures 4(c, d). Each color represents a separate evaluation depth, e.g. the rightmost green line represents the estimates for the evaluation at a 6000 nm depth, thus showing the estimates from which the bias and precision in Figures 4 (a, b) were calculated. We can observe that the cspline estimates tend to collapse to the *Z* = 0 or *Z* = −500 regions at larger depths, which results in the bias increase outside those regions, whereas the diffspline estimations remain unaltered throughout all depths.

### Interpolation evaluation

Next, we evaluated the effects of the distance between calibration beads on the resulting diffspline localization performance. For each “reference” depth *ρ* ∈ *{*1000, 4000*}* nm and for each bead distance *δ* ∈ *{*100, 300, 500, 700, 1000*}* nm, we generated PSFs at depths *ρ* −*δ, ρ, ρ* + *δ, ρ* + 2 · *δ*, thus obtaining the four required calibration PSFs. The evaluation was performed at 10 evaluation depths uniformly spaced between *ρ* and *ρ* + *δ* . For each evaluation depth, the errors between the *Z*-estimates and the ground truth were calculated. The errors were combined and depicted in the boxplot shown in Figure 4(f). We observed an increase in the mean error by 5 nm with an increase in the bead distance from 100 to 1000 nm, as well as a slight increase in the standard deviation by 4 nm.

Finally, we compared the localization bias of the cspline and diffspline calibrated on uniformly and nonuniformly spaced beads. For the uniform case, we generated calibration PSFs at depths 4000, 5000, 6000, 7000, 8000 nm, and for the non-uniform case - at depths 4500, 5600, 6366, 7277, 7877, 8400 nm. We then evaluated both splines in the depth range from the second to penultimate depth of the respective calibration PSFs. We chose five equidistant interpolation points between each consecutive pair of depths and evaluated the splines at these depths. Figures 4(g, h) show that the diffspline PSF shows a consistently low bias of approximately −5 nm and a precision of around 13 nm on average in both the uniform and non-uniform cases. The cspline estimation bias and precision both rise with an increase in the estimation depth, with the bias ranging from −125 to −100 nm and the precision rising from 15 to 42 nm in uniform case and 26 to 53 nm in non-uniform case.

### Cspline calibration at depth

The experiments above were conducted with the cspline calibrated on the coverslip for clear comparison with the methodology in the previous research^13,17^. To explore the spline differences for cspline calibration at depth, we extended the interpolation evaluation experiment shown in Figure 4 (g, h) to include the cspline models calibrated at the corresponding depth ranges: 5000, 6000, 7000 nm for uniform and 5600, 6366, 7277, 7877 for non-uniform cases. Supplementary Figure S1 shows the bias and precision for both cases in panels (a) and (b) analogously to Figure 4 (g, h), while panels (c)-(f) contain bias and precision breakdowns separately.

We can observe that the cspline estimation bias is close to zero at the calibration depths. This can also be observed in Supplementary Figure S2 which depicts the *Z*-position estimates for cspline calibrated at various depths. However, the cspline bias consistently grows with the change in depth, while diffspline bias stays constant over the depths. A similar pattern is observed for the estimation precision, where the standard deviation for the cspline troughs at the calibration depth and rises with the depth change. However, for every calibration depth, the lowest estimation precision of the cspline is higher than the precision of the diffspline. These results suggest that diffspline provides a better PSF model both at the depths, where the calibration data was present, and at the depths where it interpolated the PSF.

## Conclusion

In our work, we propose the diffspline model that enables PSF calibration from bead data at nonuniformly spaced depths. The model reduced the intensity overestimation by up to 67.8 percent points and the underestimation by up to 21.8 percent points at larger depths. This, in turn, led to a consistently low bias of −5 nm for the diffspline, and its precision closely followed the CRLB, with the differences not exceeding 4 nm across the entire *Z*-range.

The diffspline-interpolated models were shown to produce estimation results on par with the on-bead evaluation, thus enabling accurate and precise PSF calibration at arbitrary depths between those present in the bead stack. We also showed that an increase in the distance between the calibration beads influences the diffspline localization errors. The increase in the bead distance from 100 to 1000 nm induces an estimation bias of approximately 5 nm owing to interpolation errors.

Moreover, our approach reduced the computational complexity of the spline calibration from *O*(4^*d*^*V*) to *O*(*V*) as we demonstrated in Supplementary Note S1, which suggests potential applicability for higher-dimensional splines. However, one of the limitations of our current approach is the reduction in the continuity requirement from *C*^2^ in cspline to *C*^1^ in diffspline. Future research could be directed at including the second-order continuity into the Catmull-Rom spline by expanding the spline order to 5. This way, a quintic Catmull-Rom spline would include extra smoothness requirements in its definition and thus have extended interpolation capacity which could alleviate the interpolation-induced errors.

## Author contributions statement

S. K. wrote the code, conducted experiments, and wrote the manuscript. D.K. and S.H. provided feedback on the results and writing. J.C. conducted the derivations and developed the code for the 3D Catmull-Rom spline model. C.V. provided feedback on the writing. C.S.S. supervised the project.

## Additional information

## Acknowledgements

S.K., D.K., S.H., J.C., and C.S.S. were supported by the Netherlands Organisation for Scientific Research (NWO), under NWO START-UP project no. 740.018.015.

## Supporting citations

References ^31,32^ appear in the supporting information.

## Code availability

The code that supports the findings of this study is openly available on GitHub^33^ at https://github.com/qnano/diffspline.

## Data availability

The data that support the findings of this study are openly available in 4TU.ResearchData^34^ at doi.org/10.4121/5f75fadc-8f80-4e27-a049-1137031b19c0.

