## Supplementary for "Using splines for point spread function calibration at non-uniform depths in localization microscopy"

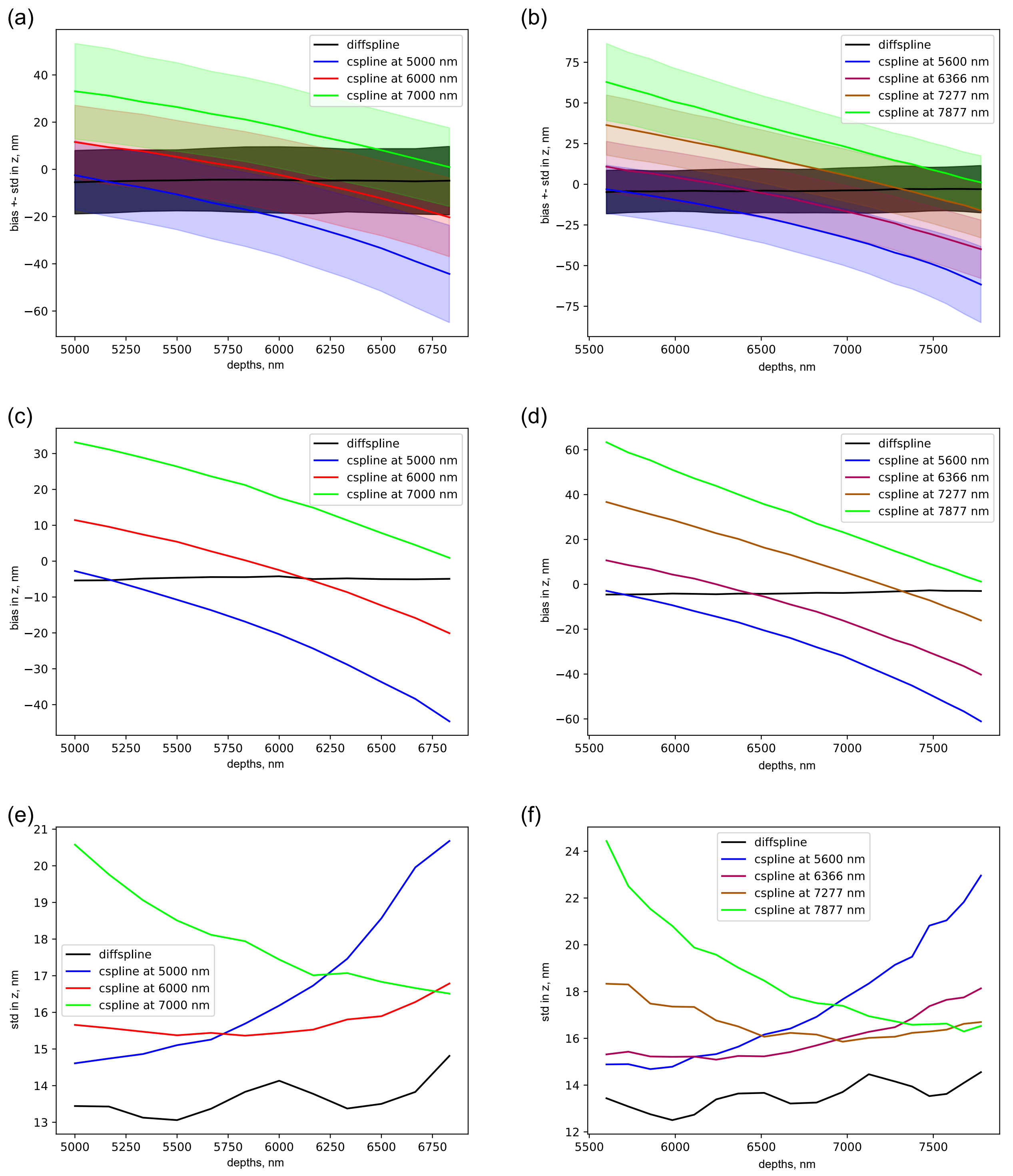


**Supplementary Figure S1.** Estimations on uniformly spaced beads at 5000,6000,7000 nm and on non-uniformly spaced beads at 5600, 6366, 7277, 7877 nm. Cspline calibrated not on the coverslip, but on uniformly and non-uniformly spaced beads directly, starts to deviate in both bias (main line) and precision (error bounds) with the increase in the distance from the calibration depth.

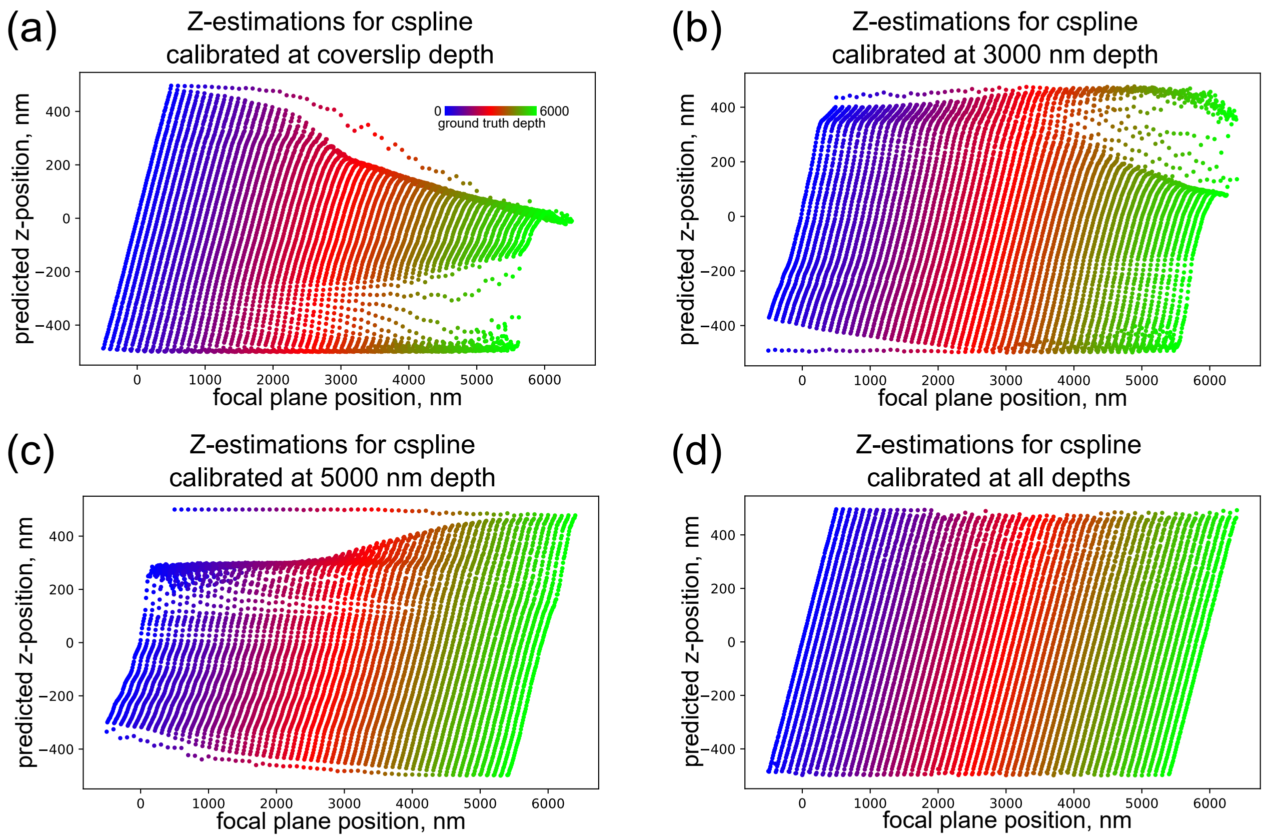


**Supplementary Figure S2.** Estimations of z-positions for cspline calibrated (a) on the coverslip, (b) at 3000 and (c) 5000 nm depths, and (d) at all depths from 0 to 6000 nm.

### Supplementary Note S1 - Spline methodology

In this note, we discuss the definitions of the spline functions we employ in our work. First, we specify the 3D cspline definition from previous work and outline the problems with extending it to 4D. Then, we provide a one-dimensional Catmull-Rom definition to exemplify how it solves the aforementioned problems the cspline has. Next, we define non-uniform version of the Catmull-Rom spline, and show how it extends to 4D splines.

#### *Cubic spline*

The 3D cubic spline (cspline) function for a PSF is implemented as a set of third-degree polynomials defined per voxel in each of the dimensions[1]. It is formulated as follows:

$$f_{i,j,k}(x,y,z) = \sum_{m=0}^{3}\sum_{n=0}^{3}\sum_{o=0}^{3} A_{i,j,k,m,n,o}\left(\frac{x-t_i}{\Delta t}\right)^m \cdot \left(\frac{y-t_j}{\Delta t}\right)^n \cdot \left(\frac{z-u_k}{\Delta u}\right)^o,$$
$$(t_i \leq x \leq t_{i+1}, t_j \leq y \leq t_{j+1}, u_k \leq z \leq u_{k+1}, \Delta t = t_{i+1}-t_i = t_{j+1}-t_j, \Delta u = u_{k+1}-u_k), \tag{1}$$

where $i$, $j$, $k$ are the voxel indices, $t_i$, $t_j$, $u_k$ are voxel coordinates, and $x$,$y$,$z$ are the evaluation coordinates in the $X$, $Y$, $Z$ dimensions respectively. The interval length, which is equivalent to the voxel size for the given dimension, is defined by $\Delta t = t_{i+1} - t_i = t_{j+1} - t_j$ in $X$ and $Y$, and by $\Delta u = u_{k+1} - u_k$ for $Z$. Finally, $A$ is the polynomial coefficient matrix of size $4^3 = 64$, where 4 is the number of spline degrees and 3 is the number of dimensions.

From this definition, it follows that the number of coefficients in the matrix $A$ to solve for depends exponentially on the number of dimensions the spline function is defined for. For instance, the 3D PSF model calibration from an image stack of size $30 \times 30 \times 80$ voxels in $XYZ$ would require solving for $4.6 \times 10^6$ coefficients. However, extending cubic splines to 4D would result in $4^4 \times m \times 80 \times 30 \times 30 = 18.4 \times m \times 10^6$ coefficients for $m$ planes in the fourth dimension. This roughly translates to 18.4 MB and $73.6 \times m$ MBs worth of memory in the float datatype. Thus, we can observe a rise in complexity with the growth of dimensions which hampers the possibilities for cspline applications for the problem at hand.

#### *Catmull-Rom spline*

The proposed diffspline model is a Catmull-Rom spline, which is in turn a specific case of a Hermite spline. Here, we describe how the Hermite spline is built. Subsequently, we show how the Catmull-Rom spline is constructed from the Hermite spline.

**Hermite spline** A Hermite spline is defined for segment $t_i, t_{i+1}$ by two points $(t_i, p_i)$, $(t_{i+1}, p_{i+1})$ and two tangents $p'_i, p'_{i+1}$ at those points. The Hermite spline is constrained to interpolate through the two points and to have the given tangents at those points:

$$\begin{bmatrix} f_i(0) \\ f_i(1) \\ f'_i(0) \\ f'_i(1) \end{bmatrix} = \begin{bmatrix} p_i \\ p_{i+1} \\ p'_i \\ p'_{i+1} \end{bmatrix}, \tag{2}$$

where $f_i(0)$ and $f_i(1)$ represent the spline function value at the beginning and the end of the given segment $t_i$, $t_{i+1}$. For a cubic Hermite spline $f_i(r) = ar^3 + br^2 + cr + d$ with $r = \frac{x-t_i}{t_{i+1}-t_i} \in [0,1]$, $t_i \leq x \leq t_{i+1}$, substitution of the polynomial results in a fully determined system of 4 equations to obtain 4 coefficients:

$$\begin{bmatrix} f_i(0) \\ f_i(1) \\ f'_i(0) \\ f'_i(1) \end{bmatrix} = \begin{bmatrix} d \\ a+b+c+d \\ c \\ 3a+2b+c \end{bmatrix} = \underbrace{\begin{bmatrix} 0 & 0 & 0 & 1 \\ 1 & 1 & 1 & 1 \\ 0 & 0 & 1 & 0 \\ 3 & 2 & 1 & 0 \end{bmatrix}}_{\mathbf{G}} \begin{bmatrix} a \\ b \\ c \\ d \end{bmatrix} = \begin{bmatrix} p_i \\ p_{i+1} \\ p'_i \\ p'_{i+1} \end{bmatrix} \tag{3}$$

To find the spline coefficients $a$, $b$, $c$ and $d$, we solve the system of equations. Using matrix inversion, we find the following solution:

$$\begin{bmatrix} a \\ b \\ c \\ d \end{bmatrix} = \mathbf{G}^{-1} \begin{bmatrix} p_i \\ p_{i+1} \\ p'_i \\ p'_{i+1} \end{bmatrix} = \begin{bmatrix} 2 & -2 & 1 & 1 \\ -3 & 3 & -2 & -1 \\ 0 & 0 & 1 & 0 \\ 1 & 0 & 0 & 0 \end{bmatrix} \begin{bmatrix} p_i \\ p_{i+1} \\ p'_i \\ p'_{i+1} \end{bmatrix} \tag{4}$$

Therefore, we obtain the final spline definition for the given segment:

$$f_i(r) = \begin{bmatrix} r^3 & r^2 & r & 1 \end{bmatrix} \begin{bmatrix} 2 & -2 & 1 & 1 \\ -3 & 3 & -2 & -1 \\ 0 & 0 & 1 & 0 \\ 1 & 0 & 0 & 0 \end{bmatrix} \begin{bmatrix} p_i \\ p_{i+1} \\ p'_i \\ p'_{i+1} \end{bmatrix}, \tag{5}$$

where $r = \frac{x-t_i}{t_{i+1}-t_i} \in [0,1]$, $t_i \leq x \leq t_{i+1}$, and $f_i(r) \in (p_i,\, p_{i+1})$.

**Catmull-Rom spline** A Catmull-Rom spline defines an interpolation function on a segment $t_i$, $t_{i+1}$ characterized by four consecutive points $p_{i-1}$,$p_i$, $p_{i+1}$,$p_{i+2}$. It substitutes the tangent values of the Hermite spline with a first order finite difference approximation of the first derivative. Specifically, it uses central difference approximation. By taking the tangent values $p'_i = \frac{p_{i+1}-p_{i-1}}{2}$, $p'_{i+1} = \frac{p_{i+2}-p_i}{2}$, we arrive to the following definition of Catmull-Rom spline:

$$f_i(r) = \begin{bmatrix} r^3 & r^2 & r & 1 \end{bmatrix} \underbrace{\begin{bmatrix} 2 & -2 & 1 & 1 \\ -3 & 3 & -2 & -1 \\ 0 & 0 & 1 & 0 \\ 1 & 0 & 0 & 0 \end{bmatrix}}_{\mathbf{H}} \underbrace{\begin{bmatrix} 0 & 1 & 0 & 0 \\ 0 & 0 & 1 & 0 \\ -0.5 & 0 & 0.5 & 0 \\ 0 & -0.5 & 0 & 0.5 \end{bmatrix}}_{\mathbf{D}} \begin{bmatrix} p_{i-1} \\ p_i \\ p_{i+1} \\ p_{i+2} \end{bmatrix}, \tag{6}$$

where $r = \frac{x-t_i}{t_{i+1}-t_i} \in [0,1]$, $t_i \leq x \leq t_{i+1}$, and $f_i(r) \in (p_i,p_{i+1})$. The matrix $\mathbf{H}$ contains the solution to the interpolation and $C^1$ continuity requirements from Equation 5, and matrix $\mathbf{D}$ shows the Catmull-Rom approximation of tangents with points.

This spline definition reduces the computational complexity compared to csplines since it requires only the pixel values for interpolation. So, in the aforementioned example image stack, the interpolation would require $4 \times 80 \times 30 \times 30$ voxel values, which equate to 1.15 MBs worth of memory. We can observe that it neither scales exponentially with the spline dimensionality, nor depends on the number of planes in the fourth dimension.

However, the Catmull-Rom tangents are devised with the assumption of uniform parameter spacing $t$. Therefore, applying this spline definition would result in highly inaccurate tangents for the spline shape.

#### *Non-uniform Catmull-Rom spline*

Barry and Goldman[2] proposed a recursive algorithm that allows to construct Catmull-Rom spline for arbitrary parameter spacing. We incorporate Barry and Goldman's definition to calculate the spline tangents in the Catmull-Rom formulation. In their definition, the finite difference for the tangents is weighted by the relative interval distance:

$$p'_i = \frac{\Delta_i(p_i - p_{i-1})}{\Delta_{i,i-1}\Delta_{i-1}} + \frac{\Delta_{i-1}(p_{i+1} - p_i)}{\Delta_{i,i-1}\Delta_i}, \tag{7}$$

where $\Delta_i = t_{i+1} - t_i$, $\Delta_{i-1} = t_i - t_{i-1}$, $\Delta_{i,i-1} = \Delta_i + \Delta_{i-1} = t_{i+1} - t_{i-1}$. The two tangents $p'_i$ and $p'_{i+1}$ for a given segment therefore take the form:

$$\begin{aligned} p'_i &= \frac{\Delta_{i-1}}{\Delta_{i,i-1}\Delta_i} p_{i+1} + \left(\frac{\Delta_i}{\Delta_{i-1}} - \frac{\Delta_{i-1}}{\Delta_i}\right)\frac{1}{\Delta_{i,i-1}} p_i - \frac{\Delta_i}{\Delta_{i,i-1}\Delta_{i-1}} p_{i-1} \\ p'_{i+1} &= \frac{\Delta_i}{\Delta_{i+1,i}\Delta_{i+1}} p_{i+2} + \left(\frac{\Delta_{i+1}}{\Delta_i} - \frac{\Delta_i}{\Delta_{i+1}}\right)\frac{1}{\Delta_{i,i+1}} p_{i+1} - \frac{\Delta_{i+1}}{\Delta_{i,i+1}\Delta_i} p_i. \end{aligned} \tag{8}$$

Expressing the tangents through the quadruplet of the points $p_{i-1}$,$p_i$,$p_{i+1}$,$p_{i+2}$, we arrive at the following general non-uniform Catmull-Rom spline definition:

$$\begin{gathered} f_i(r) = \begin{bmatrix} r^3 & r^2 & r & 1 \end{bmatrix} \underbrace{\begin{bmatrix} 2 & -2 & 1 & 1 \\ -3 & 3 & -2 & -1 \\ 0 & 0 & 1 & 0 \\ 1 & 0 & 0 & 0 \end{bmatrix}}_{\mathbf{H}} \underbrace{\begin{bmatrix} 0 & 1 & 0 & 0 \\ 0 & 0 & 1 & 0 \\ \alpha_1 & \alpha_2 & \alpha_3 & 0 \\ 0 & \beta_1 & \beta_2 & \beta_3 \end{bmatrix}}_{\mathbf{D}'} \begin{bmatrix} p_{i-1} \\ p_i \\ p_{i+1} \\ p_{i+2} \end{bmatrix}, \\ \begin{bmatrix} \alpha_1 \\ \alpha_2 \\ \alpha_3 \end{bmatrix} = \begin{bmatrix} 0 & -1 \\ -1 & 1 \\ 1 & 0 \end{bmatrix} \begin{bmatrix} \frac{\Delta_1}{\Delta_2} \\ \frac{\Delta_2}{\Delta_1} \end{bmatrix} \cdot \frac{\Delta_2}{\Delta_1 + \Delta_2}, \\ \begin{bmatrix} \beta_1 \\ \beta_2 \\ \beta_3 \end{bmatrix} = \begin{bmatrix} 0 & -1 \\ -1 & 1 \\ 1 & 0 \end{bmatrix} \begin{bmatrix} \frac{\Delta_2}{\Delta_3} \\ \frac{\Delta_3}{\Delta_2} \end{bmatrix} \cdot \frac{\Delta_2}{\Delta_2 + \Delta_3}, \end{gathered} \tag{9}$$

where $\Delta_1 = t_i - t_{i-1}, \Delta_2 = t_{i+1} - t_i, \Delta_3 = t_{i+2} - t_{i+1}$, $r = \frac{x-t_i}{\Delta_2} \in [0,1]$, $t_i \leq x \leq t_{i+1}$ and $f(x) \in (p_i, p_{i+1})$.

This adaptation incorporates the distance between parameters into its calculation of the tangents, while still keeping the Catmull-Rom structure with its complexity advantages. Note that the number of calculations required to calculate the interpolation increases compared to the uniform case (due to matrix $\mathbf{D}'$ in Equation 9 changing for every parameter interval). This can be alleviated by pre-computing the matrix for each parameter interval at the expense of memory on the same scale.

#### 4D PSF model

In order to build the 4D PSF spline model $f(x,y,z,d)$, we employ the non-uniform Catmull-Rom definition from Equation 9 to interpolate over all four dimensions.

$$
\begin{aligned}
f_{i,j,k,l}(x,y,z,d) = \sum_{m=0}^{3}\sum_{n=0}^{3}\sum_{o=0}^{3}\sum_{p=0}^{3} \left(\frac{x-t_i}{\Delta t}\right)^m \cdot \left(\frac{y-t_j}{\Delta t}\right)^n \cdot \left(\frac{z-u_k}{\Delta u}\right)^o \cdot \left(\frac{d-d_l}{\Delta d}\right)^p \\
\cdot \sum_{m'=0}^{3}\sum_{n'=0}^{3}\sum_{o'=0}^{3}\sum_{p'=0}^{3} A_{m,m'}A_{n,n'}A_{o,o'}A_{p,p'}V_{i+m'-1,j+n'-1,k+o'-1,l+p'-1}, \\
(t_i \leq x \leq t_{i+1}, t_j \leq y \leq t_{j+1}, u_k \leq z \leq u_{k+1}, \Delta t = t_{i+1} - t_i = t_{j+1} - t_j, \Delta u = u_{k+1} - u_k)
\end{aligned}
\tag{10}
$$

where $x$ and $y$ are shifts in the lateral direction, $z$ is the distance between the bead and focal plane, and $d$ is the depth between the bead and the coverslip $l$ is the depth index, $d_l$ is the depth coordinate, $d$ is the evaluation depth, $\Delta d$ corresponds to $\Delta_2$ from Equation 9, $A = \mathbf{HD}'$ from Equation 9, and $V$ is a four-dimensional matrix containing voxel values. The rest of the notation is defined as in Equation 1.

For the interpolation through depths $d$, we parameterize the spline with the respective bead depths $d_i$ and interpolate between the intensities of the same voxel positions. For interpolation in $X$, $Y$, and $Z$, we use uniform parameterization wherein $\Delta_1 = \Delta_2 = \Delta_3 = 1$, which turns the Equation 9 into Equation 6 and thus the interpolation becomes equivalent to the previous approaches[1,3].

We use the resulting 4D spline model to interpolate between the available depths $d_i$ for an arbitrary depth $d$ s.t. $d_{i-1} < d_i \leq d < d_{i+1} < d_{i+2} \forall i \in [2, N-2]$, where $N$ is he number of bead depths in the dataset. This entails that only four 3D PSFs are required for spline interpolation in the depth dimension, as opposed to the cspline approach which would require all depths for calibration.

#### Noise effects in spline calibration

The coefficients of the Catmull-Rom spline are determined via a fully determined system of equations, which has consequences for the effect of noise in the spline calibration. To show this, let

$$
\tilde{p} = \begin{bmatrix} \tilde{p}_{i-1} \\ \tilde{p}_i \\ \tilde{p}_{i+1} \\ \tilde{p}_{i+2} \end{bmatrix} \tag{11}
$$

describe the noised measurements of ground truths

$$
p = \begin{bmatrix} p_{i-1} \\ p_i \\ p_{i+1} \\ p_{i+2} \end{bmatrix} \tag{12}
$$

under arbitrary noise. As the system of equations for the spline coefficients is fully determined, the associated least-squares and maximum likelihood estimates for the spline coefficients have the same solution, namely the linear algebra solution[4–6]:

$$
\begin{bmatrix} a \\ b \\ c \\ d \end{bmatrix} = \mathbf{G}^{-1}\mathbf{D}' \begin{bmatrix} \tilde{p}_{i-1} \\ \tilde{p}_i \\ \tilde{p}_{i+1} \\ \tilde{p}_{i+2} \end{bmatrix} \tag{13}
$$

with $\mathbf{G}^{-1} = \mathbf{H}$ and $\mathbf{D}'$ defined as in Equation 9. Note that any noise added when obtaining the ground truth measurements $p$ is thus directly relayed into the spline coefficients through the matrix $\mathbf{G}^{-1}$. Specifically for the problem at hand, the ground truth support points $p$ are measured with fluorescence-induced background and Poisson shot noise, which will therefore be fitted into the Catmull-Rom spline coefficients.

To reduce the effects of noise in the spline coefficient estimation through least squares or maximum likelihood estimation, the system of spline equations needs to be overdetermined. This can be accomplished by deviating from the Hermite and Catmull-Rom formulations by imposing extra conditions on the spline, for example on the second derivative. Alternatively, maximum a posteriori estimation could be used to estimate the Catmull-Rom coefficients, in case a prior distribution on the spline coefficients is available[7]. However, due to the favourable computational nature of Catmull-Rom splines and the lack of prior information, neither approach is suited to reduce the noise in our calibration experiments.

For this reason, we need to minimize the effects of shot noise in our calibration experiments through preprocessing. First of all, the bead intensity in our simulations ranges between 5000 and 15000 signal photons. This brings the signal-to-background ratio up to the $250-750$ range, thereby minimizing the effect of background on the spline coefficients. Subsequently, we overlay multiple realizations of the same bead image, which results in averaging out the effects of shot noise when combined with the increased bead intensity. Under these preprocessing conditions, the effect of background and Poisson shot noise on the estimated spline coefficients are reduced.

Note that both steps are not straightforward to apply when calibrating on experimental data. In this case, we recommend the development of customized experimental preprocessing methods to take background and the Poissonian nature of the measurements into account. Alternatively, we would recommend opting for a non-Hermite spline where the spline coefficients are determined from an overdetermined system of equations, thereby enabling reduction of the noise through least squares, maximum likelihood estimation or (if a prior is available) maximum a posteriori estimation.
